## Supplementary information and figures S1-5 for "*Alasemenia*, the earliest ovule with three wings and without cupule"

**Fig. S1. Transverse sections of two seeds of *Alasemenia tria* gen. et sp. nov. a-k,** Serial sections of seed in Fig. 3a, b, at eleven levels and in ascending order (Slide PKUBC17913-13b, 12a, 12b, 11a, 11b, 10a, 10b, 9a, 9b, 8a, 8b). **j, k,** Arrows indicating integumentary lobes. **l-x,** Serial sections of seed in Fig. 3f, g, at thirteen levels and in ascending order (Slide PKUBC19798-11a, 9a, 9b, 8a, 8b, 7a, 7b, 6a, 6b, 5a, 5b, 4a, 4b). **c, f, i, p, t, w, x,** The same as those in Fig. 3c-e, h-k, respectively. Scale bars, 1 mm.

**Fig. S2. Transverse sections of one seed of *Alasemenia tria* gen. et sp. nov. a-q,** Serial sections of seed in Fig. 3l, m, at seventeen levels and in ascending order (Slide PKUBC17835-4b, 4a, 5b, 5a, 6b, 6a, 7b, 7a, 8b, 8a, 9b, 9a, 10b, 10a, 11b, 12a, 12b). **d, g, i, l, n,** The same as those in Fig. 3n-r, respectively. **m-q,** Arrows indicating integumentary lobes. Scale bars, 1 mm.

**Fig. S3. Transverse sections of two seeds of *Alasemenia tria* gen. et sp. nov. a-d,** Sections of seed in Fig. 3s, at four levels and in ascending order (Slide PKUBC18716-10a, 8a, 8b, 7a). **e-i,** Sections of another seed (at five levels, in ascending order) (Slide PKUBC18984-9b, 8b, 8a, 7b, 7a). **c, d,** The same as those in Fig. 3t, u, respectively. Scale bars, 1 mm.

**Fig. S4. Transverse sections of one seed of *Alasemenia tria* gen. et sp. nov. a-l,** Serial sections of seed in Fig. 3v, at twelve levels and in ascending order (Slide PKUBC20774-9a, 9b, 8a, 8b, 7a, 7b, 6a, 6b, 5a, 5b, 3a, 3b). **e, h, k, l,** The same as those in Fig. 3w-z, respectively. Scale bars, 1 mm.

**Fig. S5. Transverse sections of one seed of *Alasemenia tria* gen. et sp. nov. a-j,** Serial sections of seed in Fig. 3A, at ten levels and in ascending order (Slide PKUBC17904-6a, 6b, 5a, 5b, 4a, 4b, 3a, 3b, 2a, 2b). **d-f, h,** The same as those in Fig. 3B-E, respectively. Scale bars, 1 mm.

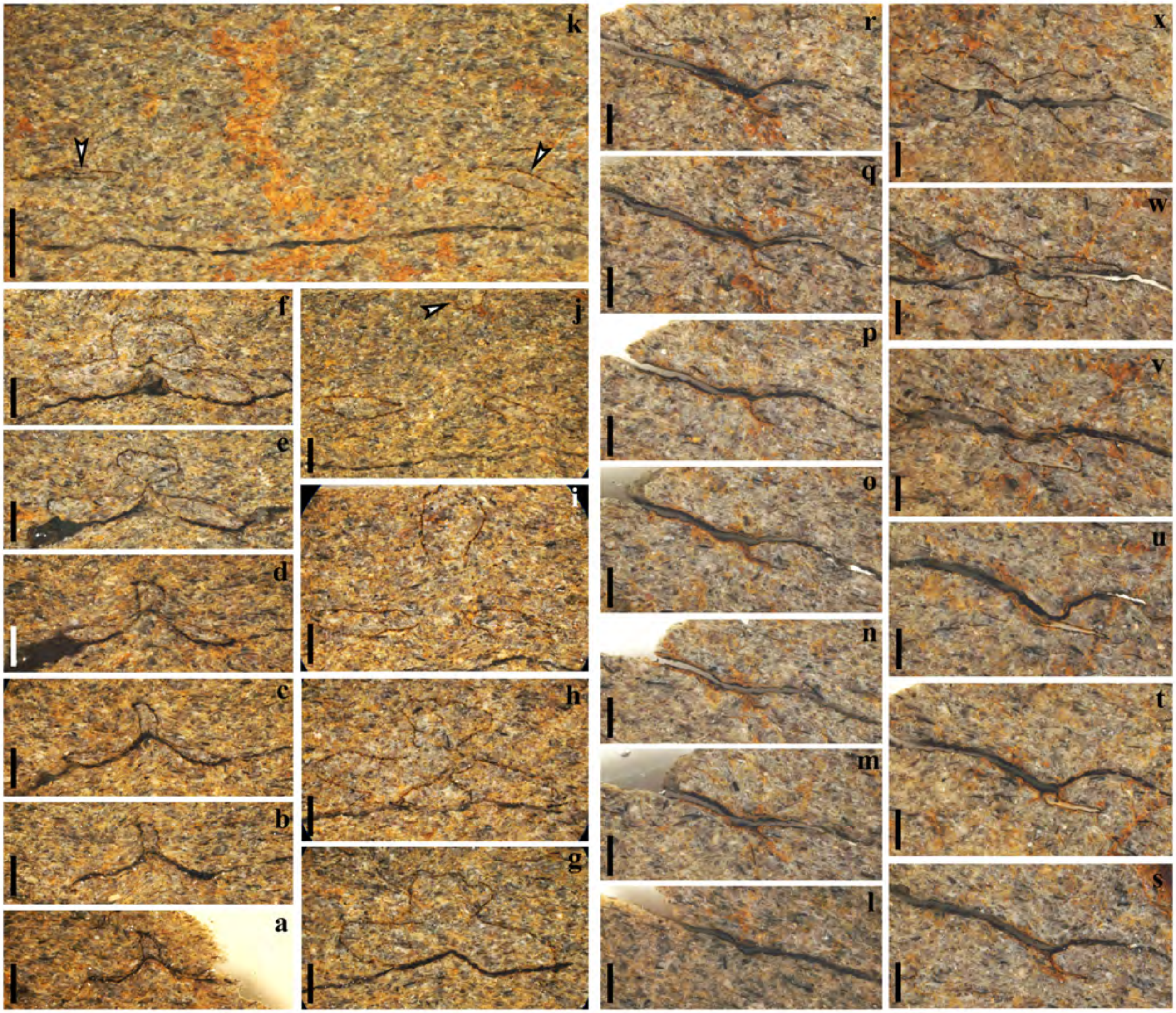


**Fig. S1**

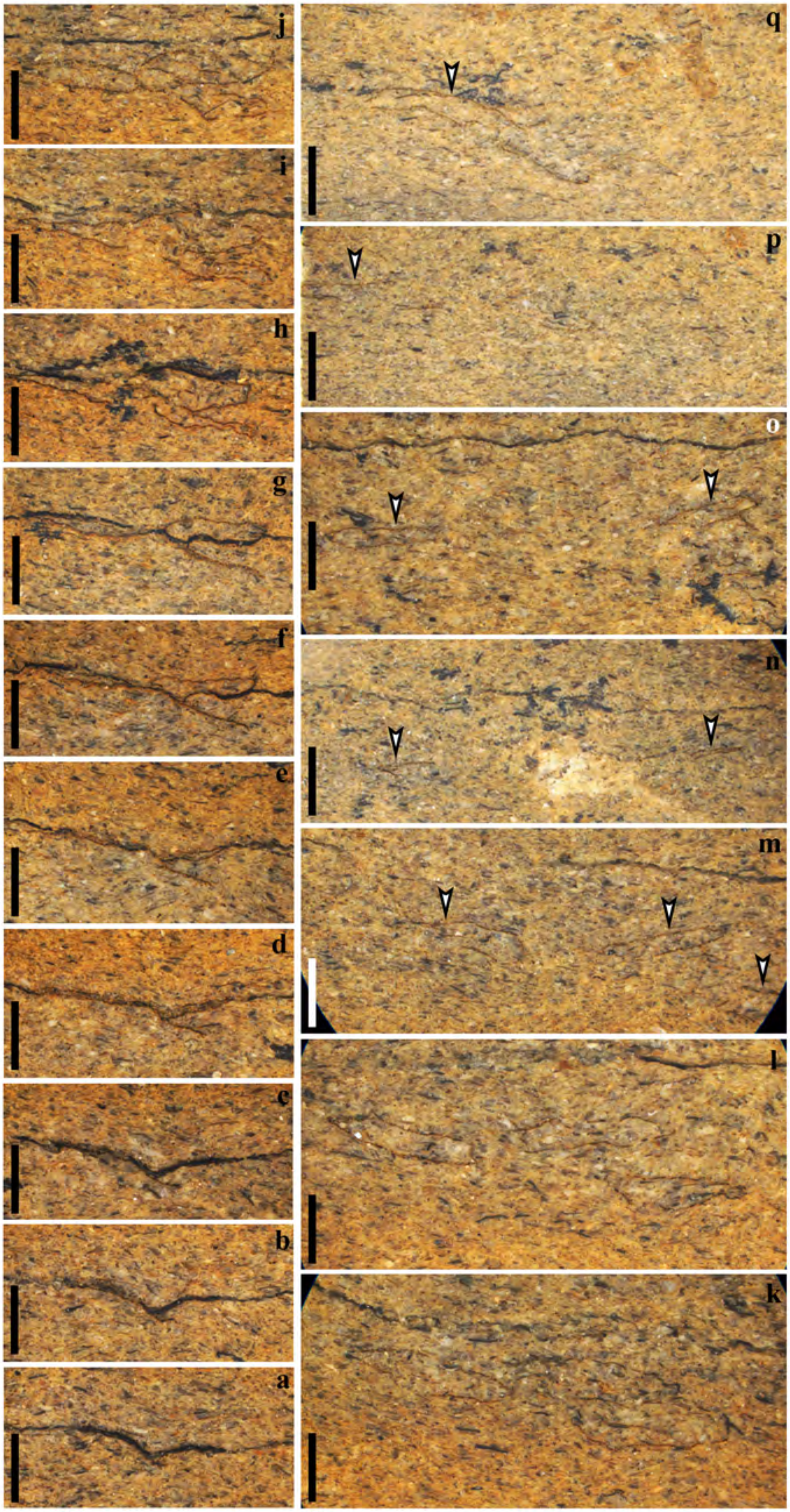


Fig. S2

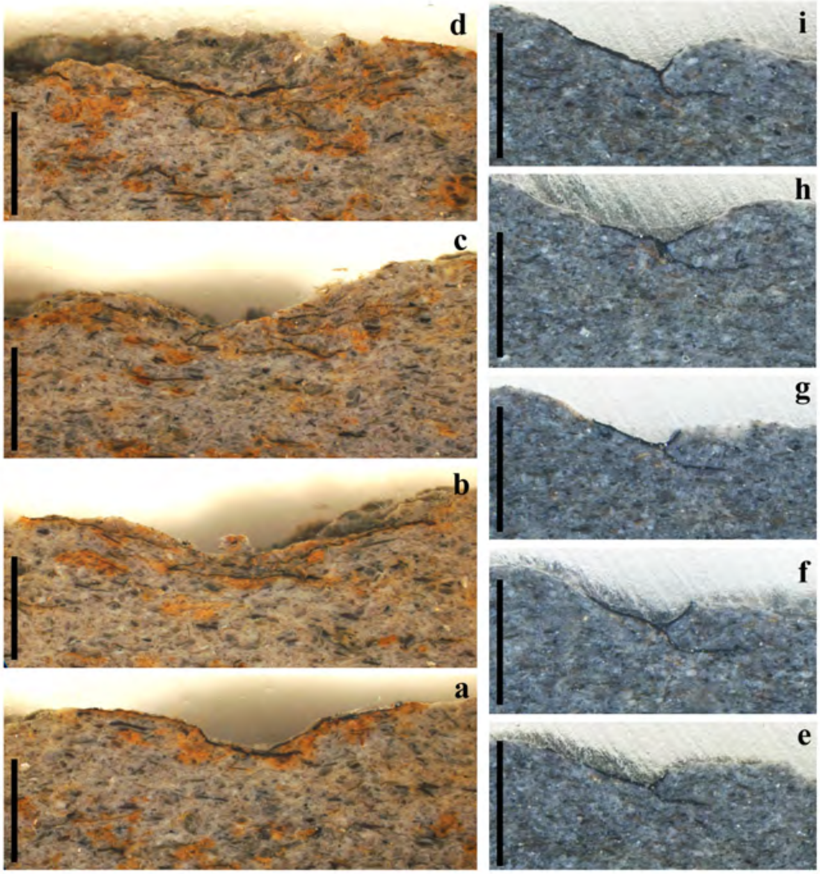


**Fig. S3**

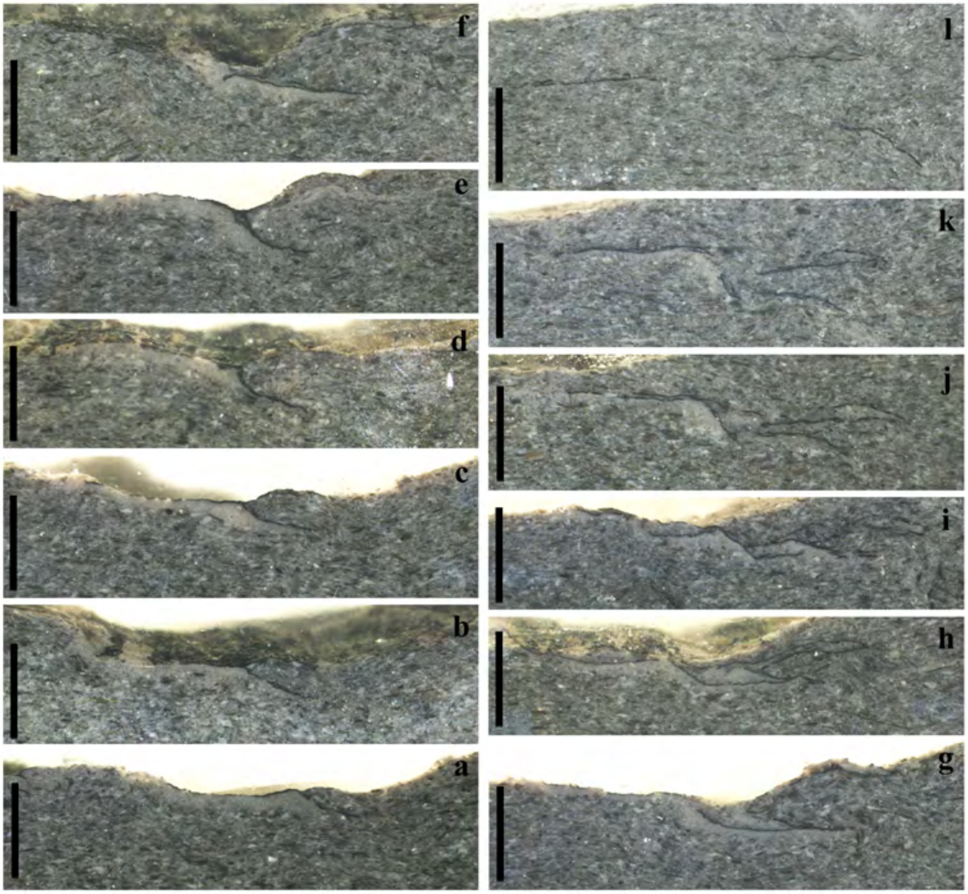


**Fig. S4**

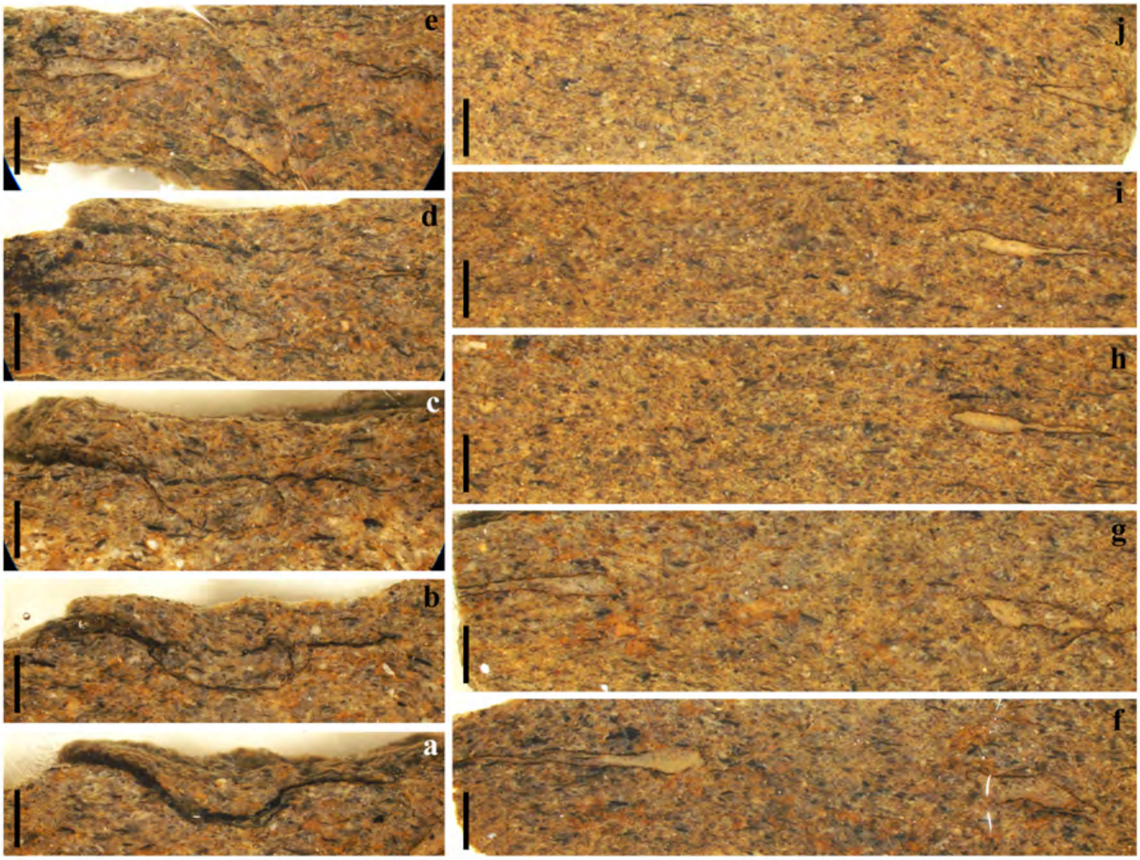


**Fig. S5**
